# The MTIST platform: a microbiome time series inference standardized test simulation, dataset, and scoring systems

**DOI:** 10.1101/2022.10.18.512783

**Authors:** Grant A. Hussey, Chenzhen Zhang, Alexis P. Sullivan, David Fenyö, Jonas Schluter

**Affiliations:** Institute for Systems Genetics, NYU Langone Health, New York, New York, United States; Department of Microbiology, NYU Langone Health, New York, New York, United States

**Keywords:** MTIST, ecological inference, microbiome, ecological interactions, ecology

## Abstract

The human gut microbiome is promising therapeutic target, but development of interventions is hampered by limited understanding of the microbial ecosystem. Therefore, recent years have seen a surge in the engineering of inference algorithms seeking to unravel rules of ecological interactions from metagenomic data. Research groups score algorithmic performance in a variety of different ways, however, there exists no unified framework to score and rank each inference approach. The machine learning field presents a useful solution to this issue: a unified set of validation data and accompanying scoring metric. Here, we present MTIST: a platform for benchmarking microbial ecosystem inference tools. We use a generalized Lotka-Volterra framework to simulate microbial abundances over time, akin to what would be obtained by quantitative metagenomic sequencing studies or lab experiments, to generate a massive *in silico* training dataset (MTIST) for algorithmic validation, as well as an “ecological sign” score (ES score) to rate them. MTIST comprises 24,570 time series of microbial abundance data packaged into 648 datasets. Together, the MTIST dataset and the ES score serve as a platform to develop and compare microbiome ecosystem inference approaches.

## 1 Introduction

The human gut microbiome comprises a diverse ecosystem of bacteria, fungi, archaea, and viruses. While the multi-kingdom diversity of the gut remain relatively unknown (Limon, Skalski, and Underhill 2017; Koskinen et al. 2017), the bacterial community continues to be the target of extensive research. At its simplest, ecological dynamics of bacterial populations colonizing the gut can be described as a dynamical system where each species interacts with itself, each other species, and the host (Foster et al. 2017; Coyte, Schluter, and Foster 2015). Out of these three interaction motifs, inter- and intra-species interactions strongly influence the overall microbiome composition (Coyte, Schluter, and Foster 2015). Specifically, elementary pairwise interactions—mutualism (+/+), competition (-/-), exploitation (+/-), commensalism (+/0), and amensalism (-/0)—form the qualitative basis of ecological dynamics (**Table 1**) (Foster and Bell 2012; Korb and Foster 2010; Mitri, Xavier, and Foster 2011; Coyte, Schluter, and Foster 2015; Xiao et al. 2017). Uncovering and mapping these interactions between bacterial taxa is critical to facilitate precision engineering of the microbiome. For example, an improved understanding could enable predictions that combat the invasion of certain human gut pathogens (Buffie et al. 2015; Morjaria et al. 2019; Taur et al. 2012) or accelerate development of “evolutionary medicine” approaches to limit pathogen growth by designing and/or introducing competitor species (Andersen et al. 2019). Additionally, the outcome of clinical interventions, like using fecal microbiota transplants (FMT) to establish a healthy microbiota, would become more predictable.

**Table 1:** Elementary pairwise interaction types between bacterial taxa.

| Interaction | Directionality | Example | Reference |
| --- | --- | --- | --- |
| Competition | -/- | the competition for a shared resource or costly microbial warfare | McNally et al. 2017; Cornforth and Foster 2013 |
| Mutualism | +/+ | the mutually beneficial exchange of nutrients and secretions reported to occur in the microbiome | Rakoff-Nahoum, Foster, and Comstock 2016; West et al. 2006; Coyte, Schluter, and Foster 2015 |
| Exploitation | +/- | where one taxon benefits to the detriment of another, which are important for reliable ecosystem assembly | Coyte et al. 2021 |
| Commensalism | +/0 | one species <u>benefits</u>, while another remains uninfluenced |  |
| Amensalism | -/0 | one species suffers, while another remains uninfluenced |  |

Numerous microbiome ecological inference algorithms have been developed to facilitate such an ecological understanding of the gut microbiome. These algorithms infer microbial interactions from time-resolved microbial abundance data, such as longitudinal 16S rRNA fecal samples (Vidanaarachchi et al. 2020; Shaw, Pao, and Wang 2016; Alshawaqfeh, Serpedin, and Younes 2017; Fisher and Mehta 2014; Gao et al. 2018; Stein et al. 2013; Buffie et al. 2015; Bucci et al. 2016; Faust et al. 2018; Rao et al. 2021; Mounier et al. 2008; Kuntal, Gadgil, and Mande 2019; Li et al. 2019; Joseph et al. 2020). Each method parameterizes the generalized Lotka-Volterra (gLV) equations, thereby uncovering which directional pairwise interaction (e.g., +/+, -/-, +/-, 0/+, and 0/-) underlies each species pair in a microbial community (**Figure 1 A**). Network-based methods, on the other hand, infer non-directional associations between species without insight on the ecological niche between the species (Kurtz et al. 2015; Nagpal et al. 2020; Dhariwal et al. 2017; Friedman and Alm 2012; Ha et al. 2020; Fang et al. 2015; Faust et al. 2012). While bacterial associations are important to answer certain scientific questions, uncovering directional interactions is imperative to engineer precision microbiota-focused therapies.

**Figure 1:**
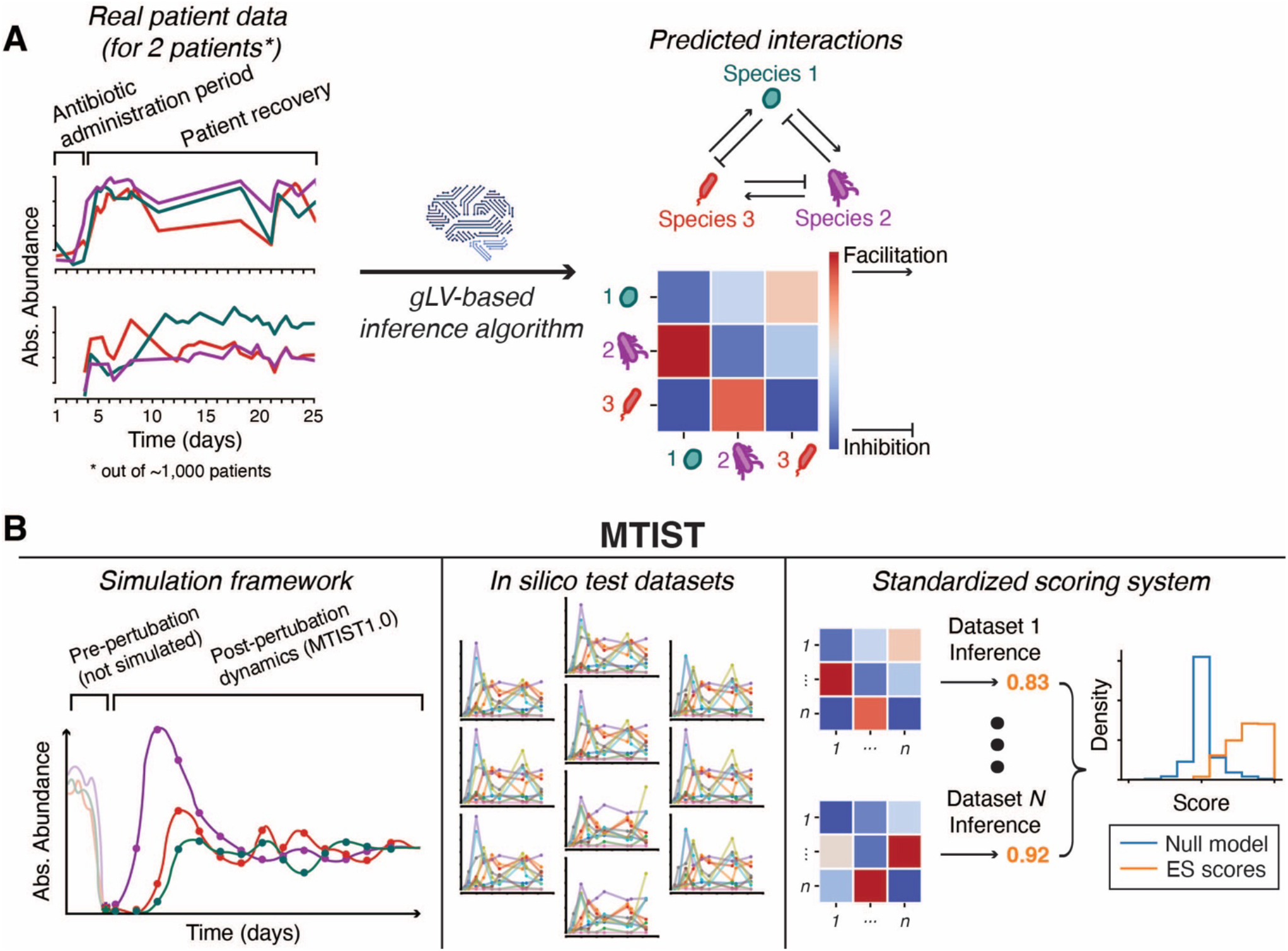
MTIST is a platform for benchmarking microbiome ecosystem inference tools. **A**. Inference algorithms learn what pairwise interactions exist between species in a bacterial community from time series data. On the left are total abundances of 3 major species measured from stool samples in two patients following strong microbiome perturbation (out of approx. 1,000, Schluter et al. 2020). Interactions can be inferred from this data using inference algorithms and are represented by the network and heatmap on the right. **B**. MTIST is made up of three parts: (1) A simulation framework, (2) a collection of *in silico* test datasets generated using that simulation framework, and (3) and scoring system to benchmark inference algorithms. The simulation framework can generate realistic *in silico* time series data inspired by patient time series like in (A). The collection of *in silico* test datasets incorporate our experience obtaining >10,000 samples from >1,000 patients. The scoring system allows one to uniformly benchmark a given tool on how well it uncovers the underlying communities that generated each *in silico* test dataset. For each of the MTIST *in silico* datasets, inference is run. An ES score is calculated per-dataset to quantify how well the inference algorithm uncovered the underlying ecosystem. All of the ES scores are then compared to a null model where inference is replaced by random chance. Inference is compared against the null model and/or to performance of other inference tools.

Properly inferring pairwise interactions is a challenge given the compositional nature of 16S rRNA sequencing (Cao et al. 2017; Aitchison 2016). Many of the gLV-based inference methods directly input 16S rRNA sequencing counts by transforming the data at each time point to relative abundances (Vidanaarachchi et al. 2020; Shaw, Pao, and Wang 2016; Alshawaqfeh, Serpedin, and Younes 2017; Fisher and Mehta 2014; Gao et al. 2018; Joseph et al. 2020; Kuntal, Gadgil, and Mande 2019). However, since the gLV model assumes that species abundances are *not* compositional but rather absolute, inference algorithms should be provided absolute cell numbers of each bacterial species whenever possible for more accurate inference (Cao et al. 2017). Indeed, several inference methods utilize 16S gene qPCR (Li et al. 2019; Stein et al. 2013; Bucci et al. 2016; Mounier et al. 2008; Rao et al. 2021; Schluter et al. 2020; Faust et al. 2018) or novel genetic “spiking” (Rao et al. 2021) to estimate absolute abundances.

Despite the diversity of inference tools, no unified method exists to assess inference algorithm performance. Instead, each research group generally uses one of two methods to analyze the inferred gLV coefficients output by their inference algorithm. First, some measure how well inferred coefficients can be used in “forward simulations” to recapitulate the original training data. Root mean squared error (Friedman and Alm 2012) or Bray-Curtis dissimilarity (Shaw, Pao, and Wang 2016; Vidanaarachchi et al. 2020) quantifies the similarity between the forward-simulated timeseries and the original training data timeseries. While such measurements provide a goodness of fit, it cannot directly conclude that an algorithm properly uncovered the pairwise interactions underlying the timeseries. A second method attempts to do just that: directly quantify how well an inference algorithm produced the exact coefficients used in generating the training data. This involves first creating an artificial microbial community parameterized in the gLV model (the “ground truth”), running numerical simulations using that ground truth to generate *in silico* training data, and attempting to infer back the exact ground truth coefficients using a prospective inference algorithm. The correctness of the inferred result is calculated by directly comparing the inferred result to the exact ground truth used to generate the training data. This method provides the most direct measure of an algorithm’s ability to uncover the interactions driving microbial time series data. Ideally, there would be known human microbiomes with well-defined microbe-to-microbe interactions, so that this method would not require the creation of artificial microbial ground truths. Since such a map does not readily exist, researchers must carefully design biologically relevant artificial microbial ecosystems. Although this method has promise, there is no current standard to benchmark an inference algorithm on *in silico* data. Here, we present MTIST, a platform for the rigorous evaluation, scoring, and comparison of microbiome ecosystem inference tools (**Figure 1 B**). Inspired by the machine learning world, we developed MTIST to benchmark algorithmic performance using a standardized methodology. This way, each tool is evaluated on identical test data and compared on equal grounds. For example, computer vision algorithms designed to read handwriting are assessed using the MNIST dataset (LeCun and Cortes 2010) to benchmark how they perform.

There are three parts of MTIST: the simulation framework, the *in silico* test dataset, and the “ecological sign” score evaluation method. The simulation framework uses the generalized Lotka-Volterra (gLV) equations and implements different sources of non-differentiable biological and technical noise caused by experimental imperfections and sample collection frequencies (**Figure 1 B, left-hand side**). The implementation of our framework was inspired by gut microbiome recovery dynamics after antibiotic perturbation observed in hospitalized cancer patients (Schluter et al. 2020), or during assembly of the infant gut microbiome (Coyte et al. 2021; Rao et al. 2021). The *in silico* test dataset generated by our simulation framework, MTIST1.0, is a collection of 648 simulated time series across biologically-relevant simulation parameters (**Figure 1 B, middle**). Finally, the benchmarking methodology scores how each new inference algorithm performs across the 648 simulated timeseries of MTIST (**Figure 1 B, right-hand side**). The evaluation methodology is designed to directly assess and compare a necessary condition for any successful ecosystem inference: how well an ecosystem inference algorithm can achieve the task of learning ecological relationships.

## 2 Methods

### 1. Building a generalized Lotka-Volterra simulation framework to generate MTIST

To generate the *in silico* datasets in MTIST, we developed a generalized Lotka-Volterra based simulation framework for microbial time series. Using our framework, we generated the datasets for 3-, 10-, and 100-species microbial communities using a wide range of simulation conditions that mimic realistic situations encountered in human microbiome time series data (**Figure 2 A-B**). In short, we used the Python programming language to simulate microbial abundances for 25 days in 100 discrete “6-hour” steps, introducing non-differentiable biological noise at the end of each step, then sub-sampled these “master time series” into individual time series to incorporate different ways microbiome studies collect stool samples.

**Figure 2:**
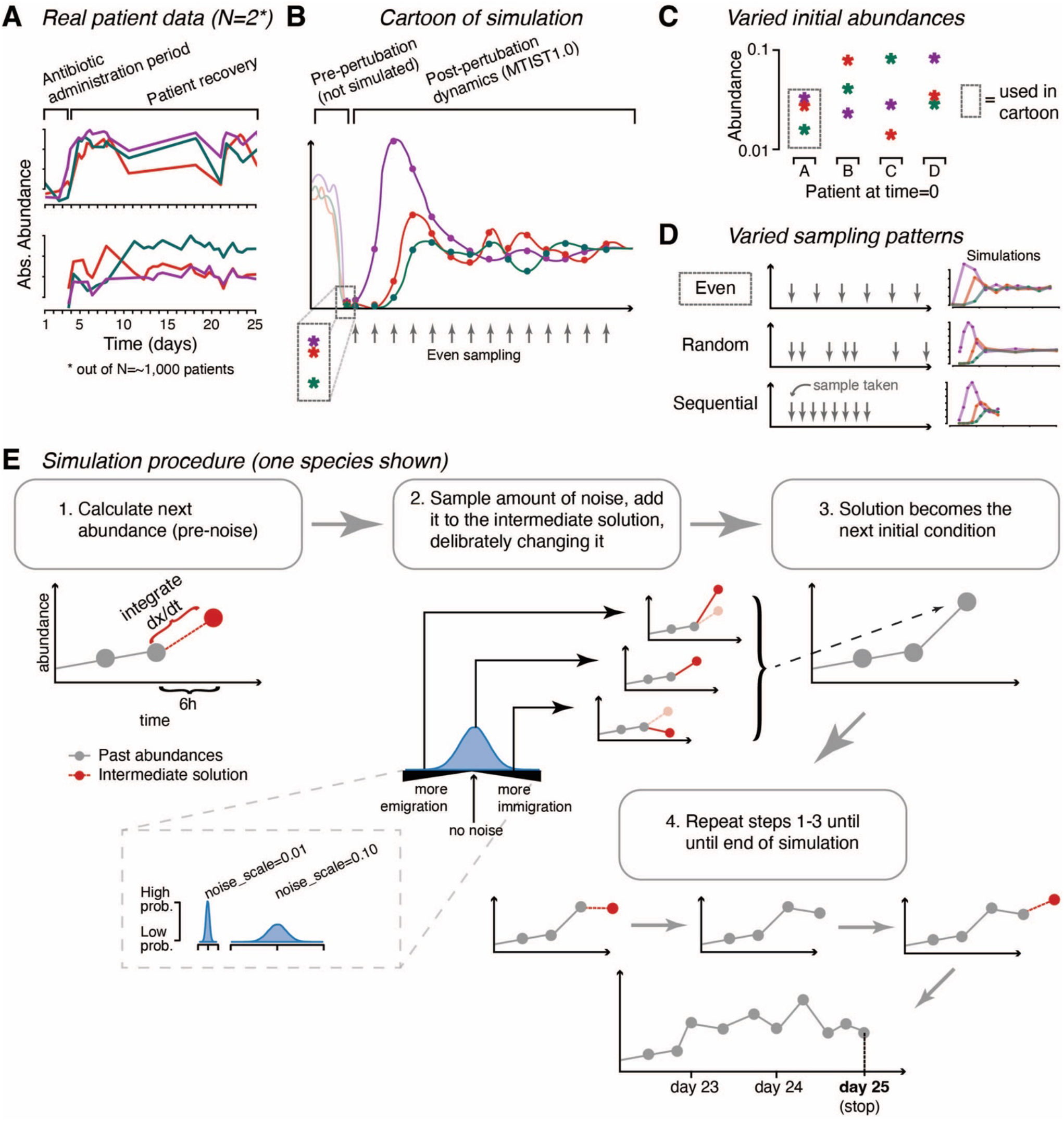
How the MTIST simulation framework works. **A**. Total abundances of 3-species measured in stool samples from two patients over time following strong microbiome perturbation (N=2 out of approx. 1,000, Schluter et al. 2020; Liao et al. 2021). **B-E** The simulation framework developed to generate MTIST1.0 **B**. Cartoon of one simulation of three species over time representative of the MTIST data. **C**. Initial species abundances vary in each simulated time series. **D**. Sampling patterns “even”, “random”, or “sequential” mimic different experimental designs or sample collection protocols as a source of real world data variability. **E**. Flowchart of our simulation procedure by example of a single species’ dynamics, displaying how we introduce changes to abundances not driven by simulated ecological interactions (noise, four times per 24h interval). The variable *noise_scale* (either 0.01 or 0.10 in MTIST1.0) alters the amount of noise added in each step and is visualized in the inset box.

For each ground truth community matrix, simulation begins by initializing each species at different low starting abundances (uniformly sampled between 0.01 and 0.1, **Figure 2 C**). This implements, for example, a perturbed microbiome state found in the gut during cancer therapy, where antibiotic and chemotherapy treatment manifests differently in each patient (Morjaria et al. 2019), or an inoculum of mixed bacterial culture in liquid media *in vitro* (Wolfe et al. 2014). After each species is initialized at low abundance, we integrate the generalized Lotka-Volterra equations (gLV) to solve for the change in species abundances at the next step using the “ode.integrate” method with the “vode” integrator from the “scipy” package (Virtanen et al. 2020). Thus, between discrete “6-hour” steps, species dynamics are governed by the deterministic gLV equations:

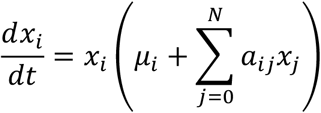

which describes 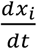, or the change in abundance of species *i*, in the 3-, 10-, or 100-species communities for each unit of time. Here, *x*_*i*_ is the abundance of species *i, µ*_*i*_ is the maximum exponential growth rate for species *i, a*_*ij*_ is the interaction coefficient between the species *i* and an interacting species *j, x*_*j*_ is the abundance of the interacting species *j*, and *N* is the total number of species present in the community. At the end of each step, we introduce non-differentiable biological noise (**Figure 2 E**), which represents stochastic migration events or changes in species abundances not driven by ecology. The noise magnitude, *n*_*i*_, for each species *i* is sampled by

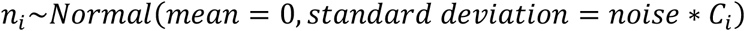

where *C*_*i*_ is the current abundance of species *i* and *noise* is a parameter dictating the scale of noise (in MTIST, either low=0.01 or high=0.10, **Figure 2 E**, see inset panel in step 2). *n*_*i*_ is then added to the corresponding species’ abundance from the intermediate solution yielded by the integration of the gLV equations over the past 6h interval (**Figure 2 E**, steps 2-3), deliberately changing the solution. The resulting abundances are then used as the initial conditions for the next 6h-integration step (**Figure 2 E**, steps 3-4). Lastly, in each step, we modeled species extinctions by setting to zero all species abundances below a very small abundance value (abundance<=0.0025).

By calculating a solution for 100 steps, we create an ideal “master time series” where stool samples are obtained from a patient exactly every 6 hours for a total of 25 days. To realistically represent a clinical or experimental microbiome sample collection protocol, we subsampled each master time series to create a set of “observed studies” to be included in our dataset. The method of sampling is dictated by one of three sampling schemes (“random”, “sequential”, or “even” sampling schemes, **Figure 2 D**) with differing numbers of samples retained (in MTIST1.0, 5 or 15). In “random”, samples are chosen at random simulated days, i.e. randomly from every fourth time point (24-hour period). In “sequential”, samples are chosen at each simulated day, i.e., at successive fourth time points, starting randomly between days 0 - 10. In “even”, samples are returned evenly throughout the entire 100 solved time points.

Figure 3 illustrates the extent of the 648 data sets comprising MTIST. For each combination of noise, sampled points, and sampling scheme, 50 unique time series were generated from the original master time series’ 50 different initial abundances for the species. These 50 unique time series were either divided into smaller sets of 5 or 25 time series or provided as a full set of 50. Such simulations were carried out for each of the 3-,10-, and 100-species communities.

**Figure 3:**
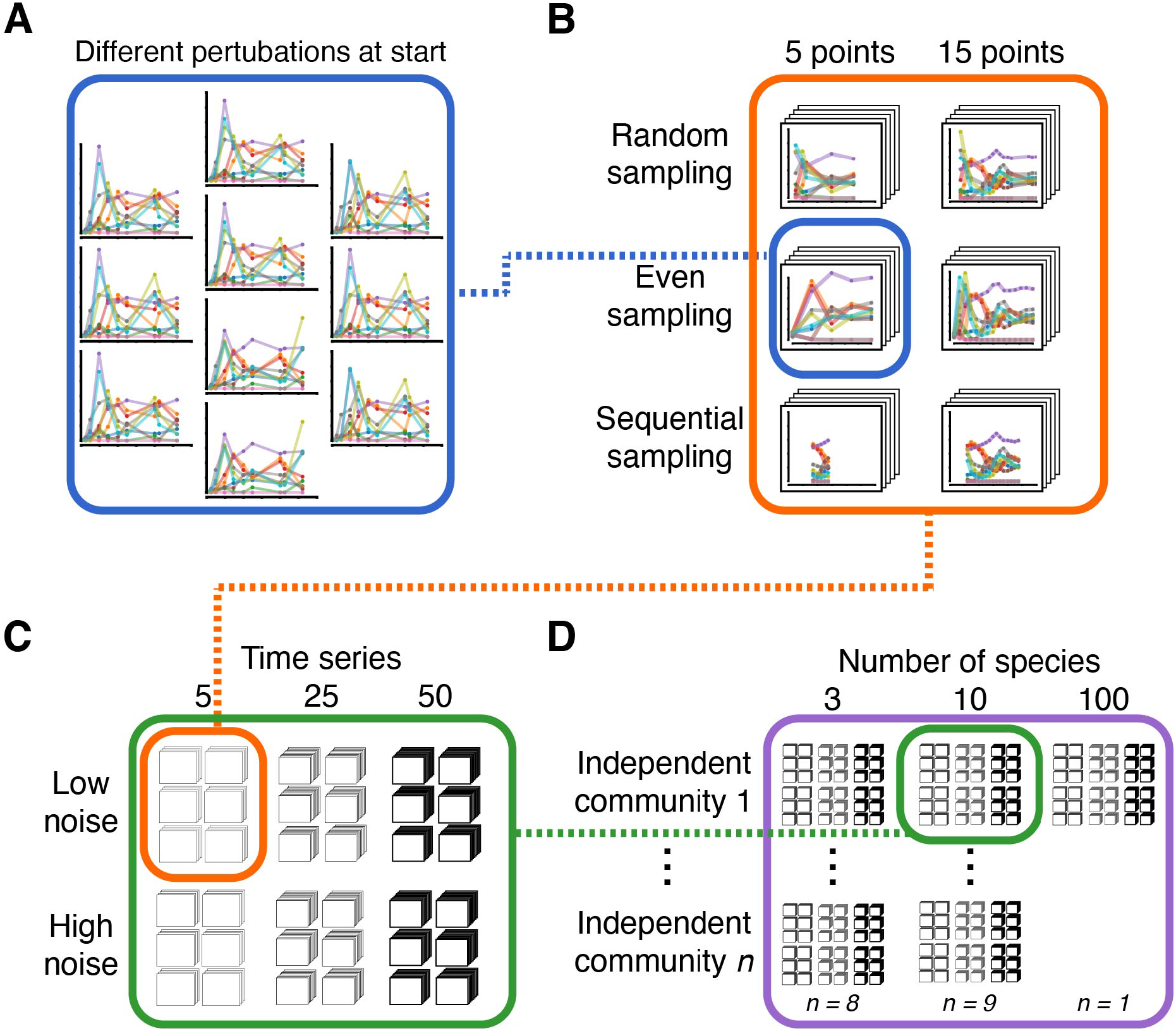
Composition of the 648 MTIST1.0 datasets. **A**. Individual time series sharing the same underlying ecosystem parameters are simulated with different initial species abundances. **B**. Using three different sampling schemes (random, even, or sequential) from a complete simulated time series at two different sampling frequencies, we mimicked sample collection variability, e.g. different experimental setups or sample collection protocols. **C** We varied the scale (low or high noise) of random abundance changes independent of ecological interactions that occur at each discrete simulation step for each simulation, as well as the number of simulated time series provided (modeling, for example, varying numbers of experiments or sampled individual hosts, i.e. 5, 25, or 50 time series). **D** The entire simulation paradigm was repeated for each for different 3- and 10-species communities as well as a single 100-species community.

The exact conditions used across the entire array of 648 datasets can be found in **Supplementary Table 1**. We provide the “mtist” package on our GitHub repository “jsevo/mtist” with examples of how all the datasets were generated.

### 2. Creating diverse synthetic ecosystems for MTIST: the coefficient matrices and maximum growth rates

For the 3-species communities, eight different interaction coefficient matrices (*A*) and maximum exponential growth rates (*μ*_i_) were used to generate the simulated time series. The 3-species community matrices were designed to encompass all key ecological interaction types: mutualism, competition, and exploitation. Since these communities are small, all species interact with each other, so there is no commensalism or amensalism. We also engineered certain inference challenges into the 3-species communities. For example, “3_sp_gt_5” and “3_sp_gt_6” share the exact same composition of interaction types: 2 exploitative relationships, and 1 mutualistic relationship. However, the interaction coefficients in each case differ so that “3_sp_gt_5” exhibits periodic behavior while “3_sp_gt_6” exhibits a more traditional growth curve behavior. Maximum exponential growth rates for these communities were calculated such that equilibrium would fall at x=1.0 for each species.

For the 10-species communities, nine different community matrices were constructed by embedding the 3-species community matrices along the diagonal of three unique, weakly-interacting 10-species “background” communities. A weakly-interacting background community was decided based on real ecosystems where most species interact rarely and weakly (McCann, Hastings, and Huxel 1998). The three weakly-interacting 10-species interaction coefficient backgrounds were generated in a two-step process. First, we imposed 82% - 93.3% of all species should interact with each other. Next, we imposed different fractions of competitive interactions (P_c_), mutualistic (P_m_) interactions, and exploitative interactions (P_e_). For the three base 10-species ecosystems prior to embedding, these values were C=91%, P_c_=12%, P_m_=5%, and P_e_=83% for base ecosystem 1 and C=73%, P_c_=18%, P_m_=0%, and P_e_=82% for ecosystems 2 and 3. Identical to the 3-species communities, maximum exponential growth rates for the 10-species communities were calculated such that equilibrium would fall at x=1.0 for each species.

The single 100-species ecosystem was generated by embedding along the diagonal three 3-species communities and three 10-species communities in a weakly-interacting 100-species background. We used the mtist function “add_n_random_species” (which imposes that interaction magnitudes for weakly-interacting species are less than half of the average interaction magnitude of the embedded community) to generate the weakly-interacting 100-species background. Growth rates for all 100 species were randomly chosen from a half-normal distribution with a scale of 0.

### 3. Implementing gLV inference for scoring methodology

Similar to most gLV inference approaches (Stein, Marks, and Sander 2015; Coyte et al. 2021), we model a gradual rate of change in species *i* abundances (*X*) over time:

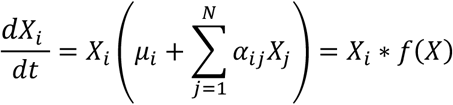

where *a*_*ij*_ represent the to be estimated pairwise interaction coefficient between species *i* and *j*. We linearize the equation,

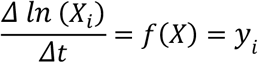

and calculate the observed log-differences, *y*_*i*_. We applied linear regression (with the LinearRegression algorithm from the scikit-learn package for the Python programming language) to estimate *y*_*i*_, using the geometric means of species abundances during Δt as predictors, 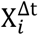, for the corresponding 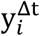, as well as an intercept. Intercept and slope coefficients therefore correspond to *µ* and *α*, respectively.

Lastly, to demonstrate that ES score differs depending on the inference method used, we repeated the above procedure but with the Ridge, RidgeCV, Lasso, LassoCV, ElasticNet, and ElasticNetCV algorithms from the scikit-learn package of the Python programming language. We also implemented MKSeqSpike (Rao et al. 2021) by adapting the author’s publicly-available code.

### 4. The “ecological sign” ES score quantifies a necessary condition for successful inference of pairwise ecological interactions

The elementary pairwise interaction types, mutualism (+/+), competition (-/-), exploitation (+/-), commensalism (+/0), and amensalism (-/0) form, to first order, the qualitative basis of ecological dynamics and are critical in understanding the underlying ecology of the microbiome (Foster and Bell 2012; Korb and Foster 2010; Mitri, Xavier, and Foster 2011; Coyte, Schluter, and Foster 2015; Xiao et al. 2017). Thus, our score incorporates the reasonable belief (Xiao et al. 2017) that inferring the sign of an interaction is the most insightful ecological metric. We present a score, the “ecological sign” ES score, to capture how well a microbiome inference algorithm performs in identifying the signs of our coefficient matrices underlying our data. The ES score therefore assesses a necessary condition for success of an inference tool: the ability to correctly infer help or harm when two species interact in a simulated world where these interactions are fundamentally true. ES is calculated by

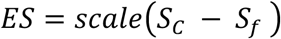

where S_c_ is the number of correct signs inferred, S_f_ is the number of incorrect opposite signs inferred. We *scale* this number to range between 0 and 1, where zero corresponds to the theoretical “worst case” scenario where all coefficients are incorrect (all positive signs are inferred as negative and vice versa) and 1 corresponds to the “ideal inference” where the exact coefficient matrices are recovered. Importantly, ES score has a few caveats. First, it applies to inference algorithms yielding pairwise interactions. Higher-order interactions that cannot be captured by gLV (Momeni, Xie, and Shou 2017; Friedman, Higgins, and Gore 2017) are not assessed. Second, the ES score can only be compared for inference results obtained over the same test data set. For this reason, we recommend annotating ES score with the data set it was calculated on (i.e., *ES*^*MTIST*1.0^). Following these instructions, inference tools based upon gLV—the most commonly assumed model for microbiome ecology (Fisher and Mehta 2014; Gao et al. 2018; Alshawaqfeh, Serpedin, and Younes 2017; Shaw, Pao, and Wang 2016; Buffie et al. 2015; Stein et al. 2013)—can be directly compared. The ES score thus assesses a necessary condition for successful ecological inference: how well an algorithm identifies ecological rules from a simulated world where those rules are exactly true.

## 3 Results

### 1. MTIST communities: all major ecological interactions are represented

MTIST is a platform for benchmarking microbial ecosystem inference algorithms. In the MTIST validation workflow, each algorithm is ranked by how well it uncovers each community matrix used to generate the MTIST datasets. For this task to be biologically relevant, we designed the bacterial communities to contain a diverse array of ecological interactions covering what is seen in a real-world human gut microbiome. In total, there are eighteen “ground truth” communities in MTIST: eight 3-species communities, nine 10-species communities, and one 100-species community. Each contain a different fraction of non-interacting pairs (0/0) and the five fundamental pairwise interactions (+/+, -/-, +/-, 0/+, and 0/-).

In the gLV-based simulations used to generate MTIST, each pairwise interaction describes how the respective species modify each other’s ecological capacity (**Figure 4 A**). For example, in an exploitative (+/-) interaction, one species benefits to the other’s detriment. Thus, the exploiter species’ ecological capacity increases while the exploited species ecological capacity decreases (**Figure 4 A**). This extends to the other fundamental interactions: in mutualism (+/+, **Figure 4 A**), both species capacities increase, while in competition (-/-, **Figure 4 A**), both species capacities decrease. Commensalism (+/0, **Figure 4 A**) and amensalism (-/0, **Figure 4 A**) respectively increase/decrease one species’ capacity while the other remains unmodified. Together, these pairwise interactions determine the evolution of species abundances over time.

**Figure 4:**
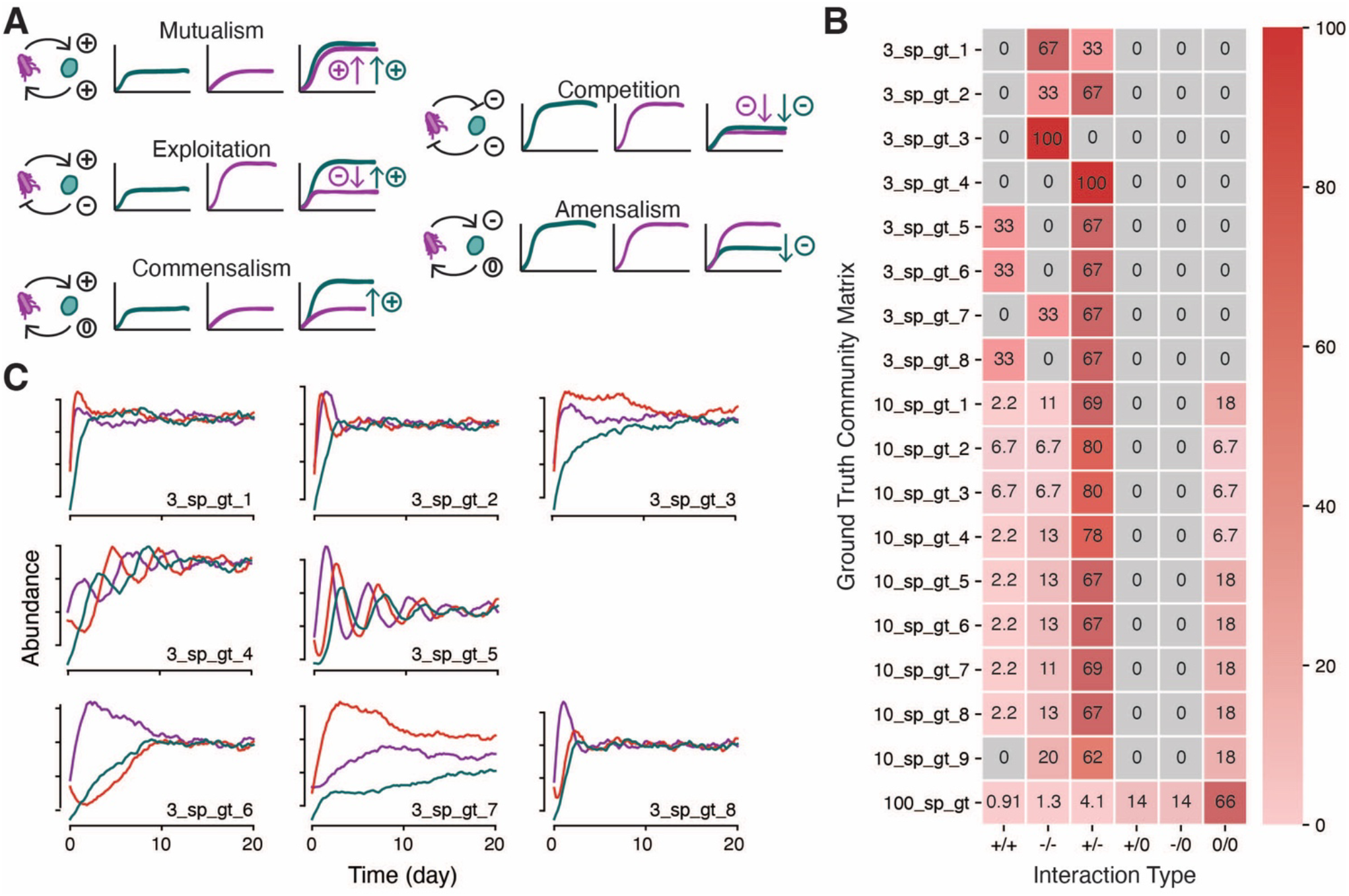
Different synthetic communities capture major types of pairwise interactions. **A**. Cartoon of the effects each interaction type has on the ecological capacity of the species involved. For example, in exploitation, the exploiting blue species reaches a higher capacity in presence of the orange species compared with growth on its own, while the exploited orange species reaches lower capacity when together. **B**. Heatmap of the percentages of certain interaction types for each ground truth community matrix. Despite having the same percentages of interaction types, different ground truths may look vastly different, as shown in **C**. Here, example time series from each 3-species community is shown. Despite “3_sp_gt_5” and “3_sp_gt_6” sharing the same composition of interaction type, the exact directionality of that interaction yields two very different simulated outcomes. This adds challenge to MTIST.

We provide examples of what the different communities look like when simulated and a summary of community matrix parameters in **Figure 4 B and C, respectively**. Briefly, in the 3-species communities, all species interact with each other, since the communities are small. Thus, mutualism (+/+), competition (-/-), and exploitation (+/-) are all present in the 3-species communities. In the 10-species communities, 6.7-18% of species do not interact with each other, whereas in the 100-species community, that number is 66%. All five interaction types (mutualism, competition, exploitation, amensalism, and commensalism) are represented.

### 2. MTIST simulations: creation of a gLV-based simulation framework incorporating real-world clinical data acquisition parameters and non-differentiable noise

The MTIST simulation framework was engineered to produce *in silico* time series consistent with our experience collecting >10,000 microbiome samples from >1,000 patients (**Figure 2 A-B**; (Schluter et al. 2020). In a clinical setting, complications with sample collection can greatly affect microbiome abundance data. To address these challenges, the MTIST simulation framework was built to generate datasets akin to an individual microbiome study. Each dataset contains a unique combination of clinical parameters: number of patients (number of time series, **Figure 2 C**), data acquisition scheme (sampling scheme, **Figure 2 D**), number of stool samples obtained (number of time points), and magnitude of biological noise (**Figure 2 E**). Benchmarking an inference algorithm on an MTIST dataset is a theoretical test to perform adequately in similar real-world study conditions.

To create the entire collection of MTIST datasets, MTIST1.0, we simulated all 3-, 10-, and 100-species communities (n=18) across biologically-relevant parameters: number of time series included (n=3), total timepoints sampled (n=2), sampling scheme (n=3), and noise scale (n=2) for a total of 648 datasets (**3**). Briefly, each individual time series begins by initializing the 3-, 10-, or 100-species microbial communities with varying low starting abundances (**Figure 2 C, Figure 3 A**) to mimic a perturbed patient microbiome or liquid medium freshly inoculated *in vitro* (**Figure 2 A**; Rao et al. 2021, Schluter et al. 2020, Coyte et al. 2021). Then, our simulation framework calculates microbial abundances in 100 discrete steps to obtain an ideal “master time series” where stool samples are taken from a patient every 4 hours. Since a realistic patient time series is far from this 100 time point ideal, we then sample either 5 or 15 time points from the master time series (**Figure 3 B**). Sampling is done in one of three different manners to create realistic, varied data representing different clinical sample collection protocols (**Figure 2 D, Figure 3 B**, see Methods). Additionally, we implement non-differentiable noise events in between each simulated time point in the master time series. This captures random species migration events or other stochastic processes that alter population densities independently from ecosystems dynamics (**Figure 2 E**). We vary the degree of these perturbation events by considering two “noise” magnitudes proportional to the current population densities (**Figure 2 E, bottom**; **Figure 3 C**). After all time series are generated, we packaged 10, 25, or 50 individual time series at each of these varied conditions into the different MTIST datasets (e.g. N=10, 25, or 50 patients enrolled in a study, **Figure 3 C**). A full list of conditions used to generate each MTIST dataset can be found in **Supplementary Table 1**.

### 3. MTIST scoring system: Algorithmic performance can be benchmarked

MTIST is a platform to determine which inference algorithms best uncover the microbial interactions that drive microbiome time series dynamics. We developed the ecological sign (ES) score to facilitate this process. ES score quantifies how well an algorithm correctly identifies the *signs* of interaction coefficients. Those signs designate which elementary pairwise interaction (mutualism, competition, exploitation, amensalism, or commensalism) exists between each pair of species and is key to facilitating precision engineering of the microbiome (Xiao et al. 2017). **Figure 5** demonstrates how two algorithms (ElasticNet regression and MKSeqSpike from Rao et al. 2021) compare in their ability to infer a single MTIST timeseries (dataset ID **125**) as quantified by ES score. Briefly, to calculate ES score, we compare the ground truth community matrix for MTIST dataset **125** (**Figure 5 A**) to the inference result of ElasticNet and MKSeqSpike (**Figure 5 B** and **C**, respectively).

**Figure 5:**
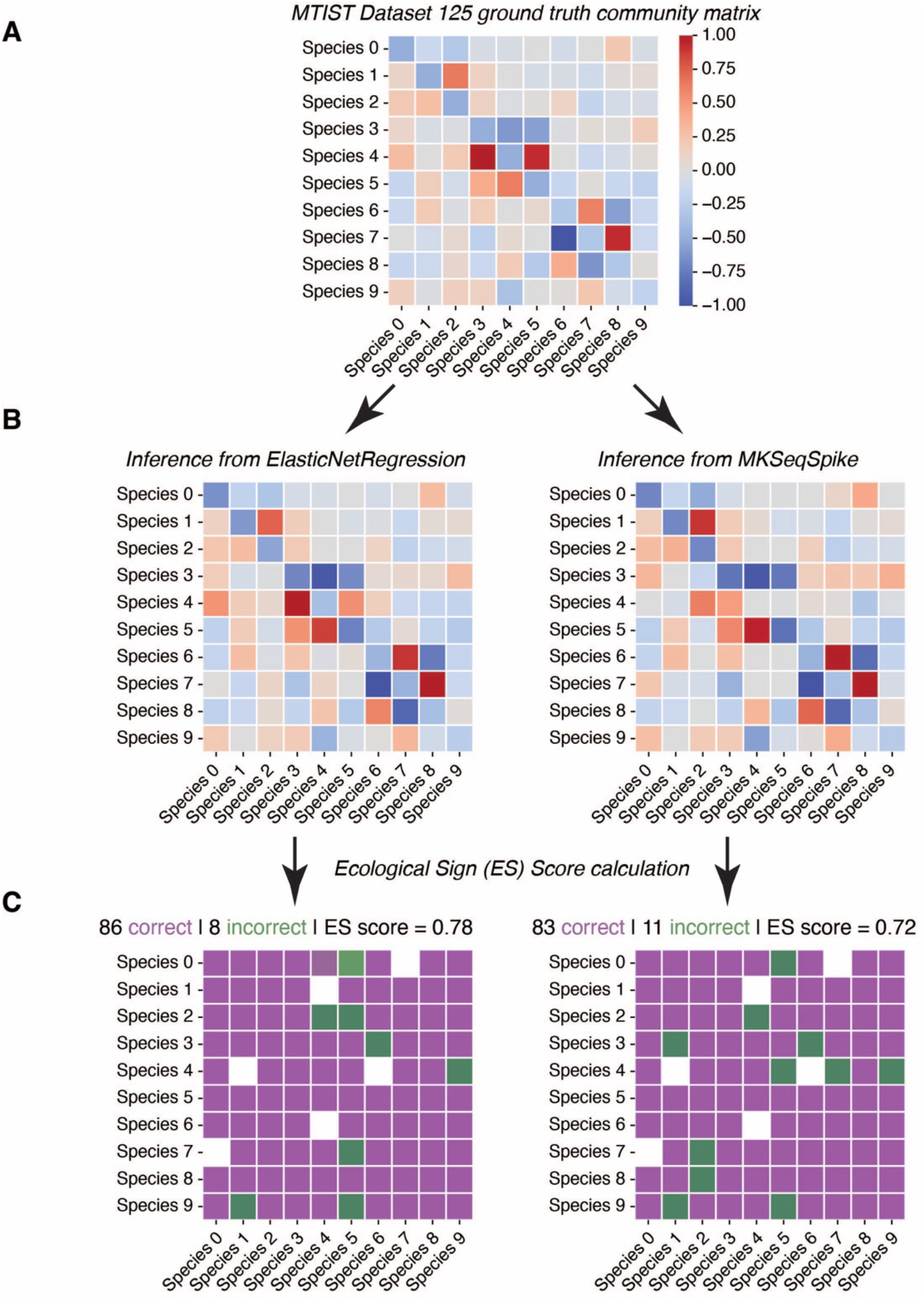
ES score can compare algorithmic performance for a single MTIST dataset. **A**. Ground truth community matrix for the 10-species ground truth from dataset ID 125. **B**. Inference result from ElasticNetRegression and MKSeqSpike tasked with inferring the interaction coefficients from MTIST dataset ID 125. **C**. For each inference result, the number of correct and incorrect sign inferences were tallied and ES score was calculated. Briefly, an inference method receives 1 point for each correct sign inference (middle column, purple “1’s”) or loses 1 point for incorrect sign inference (middle column, green “-1’s”). No penalty is given if an inference algorithm assigns “no-interaction” (i.e. a coefficient of zero). By counting the points and scaling the sum between 0 and 1 (see methods), ES score is calculated. ES score of 1 indicates perfect inference. Since the ES score of ElasticNet is greater, due to 3 more net correct sign inferences, ElasticNet can be said to perform more accurate inference for this specific dataset with its unique set of conditions.

MKSeqSpike outperforms ElasticNet since there are 10 more net correct inferences using MKSeqSpike than ElasticNet, resulting in a 0.06 point difference in ES score (0.78 vs 0.72). This demonstrates how ES score can summarize algorithmic performance for a single MTIST dataset. However, to properly capture the full range of algorithmic performance when challenged with a diversity of training data, ES score must be calculated across multiple MTIST datasets.

Each of the 648 MTIST datasets are generated with a unique combination of simulation parameters (sampling frequency, simulation noise, specific bacterial community ground truth, etc., full details for every MTIST dataset in **Supplementary Table 1**). Thus, calculating ES score for all 648 datasets provides a complete picture of algorithmic performance. **Figure 6** demonstrates how LinearRegression, RidgeRegression, ElasticNet, and MKSeqSpike inference methods perform across this MTIST subset. ES score of inference results (orange histograms) are compared to ES scores from a null model (blue histogram) where inference is replaced by random sampling from a uniform distribution as described in methods. For the 3- and 10-species communities (**Figure 6 A** and **B**, respectively), each inference algorithm generally outperforms the null model, showing that they all can appropriately learn microbial interactions from the MTIST datasets. In the 3-species case, all inference methods LinearRegression, RidgeRegression, and ElasticNetRegression achieved median ES scores of 1.0 (perfect sign inference. In the 10-species case, LinearRegression, RidgeRegression, and ElasticNetRegression each achieved comparable median ES scores of 0.78, 0.78, 0.71, respectively, while MKSeqSpike achieved a higher median ES score of 0.81. For the 100-species communities, however, no inference method outperformed the null model (**Figure 6 C**).

**Figure 6:**
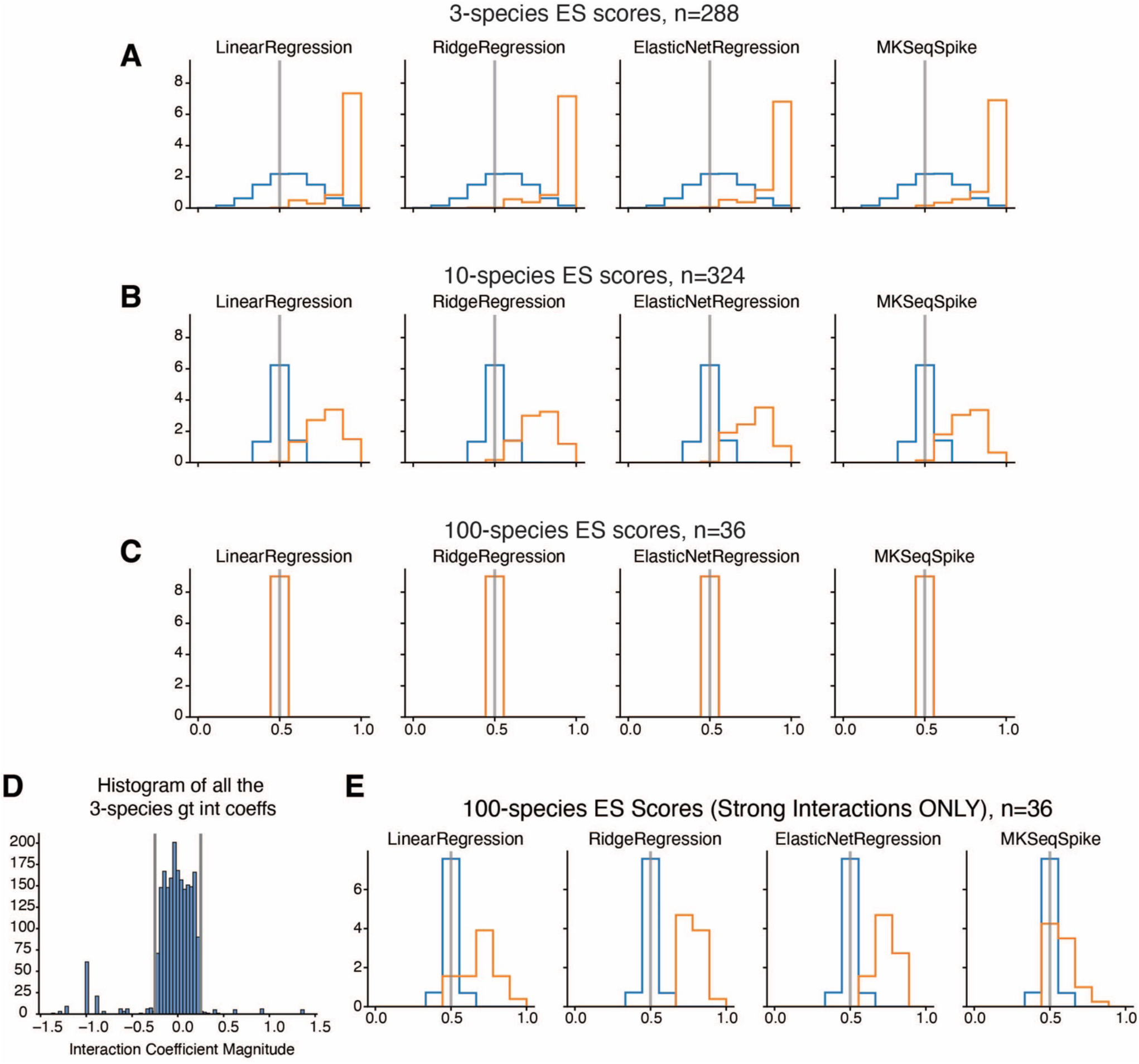
ES score across a curated MTIST subset provides a full benchmarking of inference algorithms. Histograms of ES scores across the curated MTIST subset is sufficient to benchmark inference algorithms for the (**A**) 3-species communities, (**B**) 10-species communities, and (**C**) 100-species community. Orange histograms are actual ES scores calculated once per inference method (LinearRegression, RidgeRegression, ElasticNetRegression, or MKSeqSpike) per MTIST dataset within the curated MTIST subset. Blue histograms were made from 10,000 ES scores from a null model where assignment of inferred coefficients was replaced with randomly sampling a uniform distribution (i.e., random inference). (**D**) Histogram of all interaction coefficients that comprise the 100-species ground truth community matrix. Vertical gray lines designate where the interaction coefficient magnitude is at -0.25 and 0.25. Greater than 98% of all interaction coefficients are greater than -0.25 or less than 0.25. The other ∼2% of interactions are “strong”, meaning they’re less than -0.25 or greater than 0.25 in magnitude. (**E**) Orange histograms are ES scores re-calculated only considering strongly-interacting interaction coefficients. Blue histograms are 10,000 ES scores calculated from a null model where inference was replaced by randomly sampling a uniform distribution (i.e., random inference). RidgeRegression achieves the highest median ES score.

Next, we investigated why inference with any algorithm was so poor in the 100-species communities. Since every method equally struggled on the 100-species community (median ES scores of 0.51, 0.52, 0.51, and 0.50 for LinearRegression, RidgeRegression, ElasticNetRegression, and MKSeqSpike respectively), we suspected the difficulty was due to the large number of “weakly-interacting” species present in the 100-species community. Indeed, >98% of the interaction coefficients in the 100-species ground truth community matrix fall between -0.25 < *a*_*ij*_ < 0.25 (**Figure 6 D**) compared to ∼69% and ∼4% in the 10- and 3-species case, respectively. Thus, we considered a modified ES score where correct sign inference is only considered for “strongly-interacting” species (i.e., where -0.25 > *a*_*ij*_ > 0.25). Plotting a histogram of those strong-only adjusted ES scores in **Figure 6 E**, RidgeRegression outperforms all other inference methods. For LinearRegression, RidgeRegression, ElasticNetRegression, and MKSeqSpike, the median strong-interaction-only ES score was 0.71, 0.76, 0.72, and 0.60, respectively. Thus, RidgeRegression is likely the best algorithm to properly infer interactions in bacterial communities that are 100 species or larger.

## 4 Discussion

We have presented a set of *in silico* time series data and a corresponding evaluation score, the ecological sign (ES) score, to be used by microbial ecosystem inference algorithms evaluated on the 648 datasets compiled for MTIST. We demonstrated the utility and application of this resource by evaluating a simple implementation of gLV inference. Using our ES score, we demonstrated that this simple inference approach already achieved significant learning of ecosystem rules. In particular, the large number of different datasets allowed us to investigate where this simple algorithm failed, and how especially high-temporal resolution sampling of individual timeseries supports ecological inference—with implications for microbiome study protocols.

Our standardized methodology will allow scoring and comparing new, more sophisticated tools to uncover pairwise species interactions towards an improved understanding of the microbial ecosystem in the human gut. Our simulation framework and the ES score are opinionated and focused on simple, first-order interactions between microbes. Therefore, there are several limitations to our approach. In focusing on pairwise interactions, we are excluding from our simulated ecosystems scenarios where presence of a third species might alter how a focal pair of species interact (Gonze et al. 2018; Friedman, Higgins, and Gore 2017). Furthermore, while our algorithm includes migration in the form of our noise implementation, we do not simulate (1) interactions between species that might change stochastically, over time or with changing environmental conditions, (2) interactions that might saturate, and (3) specific dynamics arising from chemical resource mediated interactions (Gonze et al. 2018; Momeni, Xie, and Shou 2017). For the latter case, we provide an example how this may be generated, expanding MTIST beyond gLV. However, we focused on the gLV approach for this first instance of MTIST because it is the most widely applied framework for inference from human data (Bashan et al. 2016; Gonze et al. 2018; Stein et al. 2013; Buffie et al. 2015; Bucci et al. 2016; Fisher and Mehta 2014; Rao et al. 2021; Schluter et al. 2020; Coyte et al. 2021). As microbiome data become more sophisticated, including via improved quantitative metabolomics, other microbiome ecosystem inferences may be enabled. Indeed, therefore, the MTIST platform as presented herein can be expanded accordingly in future iterations.

## 6 Author Contributions

GAH and JS conceived of the analyses. JS designed the algorithm. GAH generated the MTIST data set. CZ implemented and generated the resource-mediated ecosystem work. GAH, JS, APS designed and produced the figures. GAH and JS wrote the manuscript. All authors commented on the manuscript.

## 8 Acknowledgments

We thank Sara Mitri for helpful discussions on the implementation during the early stages.

## 10 Data Availability

The MTIST dataset from this study can be found in the mtist repository on GitHub (https://github.com/jsevo/mtist). The code to generate our data is included in the same repository, together with examples in the form of Jupyter notebooks.

